# Live monitoring of ROS-induced cytosolic redox changes with roGFP2-based sensors in plants

**DOI:** 10.1101/2020.12.21.423768

**Authors:** José Manuel Ugalde, Lara Fecker, Markus Schwarzländer, Stefanie J. Müller-Schüssele, Andreas J. Meyer

**Affiliations:** Institute of Crop Science and Resource Conservation (INRES), Rheinische Friedrich-Wilhelms-Universität Bonn, Friedrich-Ebert-Allee 144, D-53113 Bonn, Germany; Institute of Plant Biology and Biotechnology, Westfälische Wilhelms-Universität Münster, Schlossplatz 8, D-48143 Münster, Germany

**Keywords:** Grx1-roGFP2, roGFP2-Orp1, ROS, plate reader, CLSM, glutathione reductase, NADPH oxidase, flg22

## Abstract

Plant cells produce reactive oxygen species (ROS) as by-products of oxygen metabolism and for signal transduction. Depending on their concentration and their site of production, ROS can cause oxidative damage within the cell, and must be effectively scavenged. Detoxification of the most stable ROS, hydrogen peroxide (H_2_O_2_), via the glutathione-ascorbate pathway may transiently alter the glutathione redox potential (*E*_GSH_). Changes in *E*_GSH_ can thus be considered as a proxy of the oxidative load in the cell. Genetically encoded probes based on roGFP2 enable extended opportunities for *in vivo* monitoring of H_2_O_2_ and *E*_GSH_ dynamics. Here, we report detailed protocols for live monitoring of both parameters in the cytosol with the probes Grx1-roGFP2 for *E*_GSH_ and roGFP2-Orp1 for H_2_O_2_, respectively. The protocols have been adapted for live cell imaging with high lateral resolution on a confocal microscope and for multiparallel measurements in whole organs or intact seedlings in a fluorescence microplate reader. Elicitor-induced ROS generation is used as an example for illustration of the opportunities for dynamic ROS measurements that can easily be transferred to other scientific questions and model systems.

## 1 Introduction

Reactive oxygen species (ROS) are formed ubiquitously in cells exposed to molecular oxygen. Superoxide (O_2_**·**^-^) is generated as a by-product of oxygenic photosynthesis, by the mitochondrial electron transport chain and extracellularly by plasma membrane-localized NADPH oxidases (RBOH) [1, 2]. Due to its extremely high reactivity with itself, other radicals and transition metals in aqueous medium, O_2_**·**^-^ has a half-life of only 1 µs in a cellular environment. It is usually rapidly converted to hydrogen peroxide (H_2_O_2_), which has a half-life of 1 ms in the cell [3]. Cellular superoxide dismutases (SODs) mediate this conversion speeding up the reaction by 10^4^ orders of magnitude compared to spontaneous conversion [4, 5].

In plants, increased production of ROS has been observed under several stress situations [6] as well as during developmental processes, such as for instance polarized growth in root hairs and pollen tubes [7, 8] or the transition from proliferation to differentiation in root tips [9]. Whilst still being reactive and potentially damaging, H_2_O_2_ also acts as a signaling molecule that induces posttranslational modifications on proteins with redox-reactive residues. These modifications may alter the structural properties and activities of proteins [10, 11]. Indirect H_2_O_2_-dependent signaling may also occur via the cellular glutathione redox buffer if significant fluxes of H_2_O_2_ are detoxified via the glutathione-ascorbate cycle, which can lead to a transient change in the glutathione redox potential (*E*_GSH_) [12, 13]. The signaling function of H_2_O_2_ implies dynamic changes in H_2_O_2_ fluxes to ensure that activation or inactivation of downstream target proteins is only transient. Most assays for H_2_O_2_ based on chemical dyes such as 2’,7’-dichlorodihydrofluorescein diacetate or dihydrorhodamine 123 can, however, only monitor ROS accumulation but fail to report any dynamics.

Limitations in the dynamic monitoring of ROS have been overcome with the development of a whole series of genetically encoded redox biosensors. Redox-sensitive GFP (roGFP) in conjunction with glutaredoxin (Grx) as a thiol-disulfide switch operator equilibrates with the local *E*_GSH_ in a reversible manner and provides dynamic information about the *E*_GSH_ in its direct vicinity, i.e. typically a specific cell compartment [14, 15]. roGFP probes have two excitation peaks with maxima at 395 nm for the protonated neutral form of the chromophore (A-band) and at 490 nm for the de-protonated anionic form (B-band) (**Fig. 1**) [16]. The two excitation peaks show opposite redox-dependent changes in fluorescence intensity and are separated by the redox-indifferent isosbestic point. With these spectral characteristics, roGFP-probes are bona fide ratiometric sensors.

**Fig 1.**
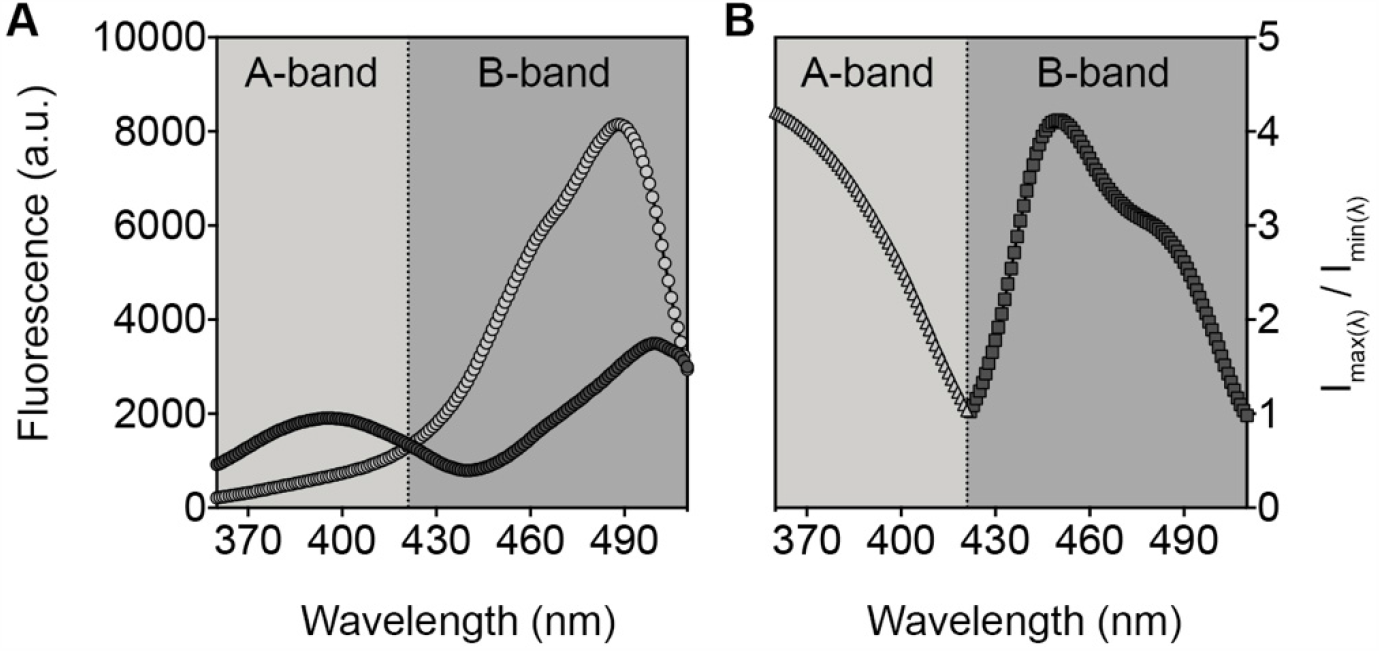
Spectral properties of purified roGFP2-Orp1. (**A**) Fluorescence excitation spectra recorded for fully reduced (light grey) or fully oxidized (dark grey) probes. Superposition of excitation spectra for reduced and oxidized sensors reveals the redox-indifferent isosbestic point at 422 nm. The two spectral areas left and right of the isosbestic point are labeled as A-band and B-band. (**B**) Relative redox-dependent changes in fluorescence intensities along the entire excitation spectrum. Fold changes in fluorescence were calculated for each wavelength from intensities in panel (A) as I_max(λ)_/I_min(λ)_, which is I_H2O2_/I_DTT_ for the A-band (light grey triangles) and I_DTT_/I_H2O2_ for the B-band (dark grey squares). Note that for the total spectral dynamic range of the sensor the relative changes at the chosen excitation wavelengths of the A- and the B-band need to be multiplied. Data are means of 4 replicates and are taken from an experimental dataset previously reported in [20], Fig. 2a.

The H_2_O_2_ sensors of the HyPer family contain the redox-active domain from the bacterial oxidation-sensitive transcription factor OxyR as redox-reactive specificity unit, which has been separated into two parts and fused to the N- and C-terminal ends of circularly permuted YFP (cpYFP). Oxidation of the reactive Cys residues of the OxyR domain by H_2_O_2_ triggers a conformational change that modifies the structural environment of the chromophore. As a result, the HyPer probes can report changes in H_2_O_2_ concentrations [17–19]. The ratiometric redox response of all these reporters enable normalized measurements that are independent of the amount of sensor protein present in cells or subcellular compartments. To respond dynamically, the sensors require re-reduction. Electrons must be provided by endogenous redox systems of the local cell compartment, such as the glutathione system or thioredoxins (TRXs), depending on the sensor [20, 21]. Most HyPer probes are extremely pH-sensitive and thus require careful pH control or parallel recording of pH [20]. This persisting drawback has been overcome only recently with the introduction of HyPer7 [19]. Another possibility for pH-independent sensing of H_2_O_2_ levels inside cells has been reported with roGFP2-based probes that use the yeast glutathione peroxidase Orp1 (also named Gpx3) or a single-cysteine variant of the peroxiredoxin Tsa2 as redox-reactive specificity domains [15, 22]. The dynamic responses of both HyPer family proteins and roGFP2-Orp1 depend on the relative rates of oxidation by H_2_O_2_ versus the rate of reduction by the interacting endogenous thiol system. Consequently, calibrations to determine absolute H_2_O_2_ levels do not carry mechanistic meaning; instead the sensors are well suited to monitor physiologically-meaningful relative redox changes in time and space.

Visualization of sensor redox status can be achieved by confocal laser scanning microscopy (CLSM) [23]. If a focus on a specific tissue area is not required or even unfavorable (e.g., due to heterogeneity), it is also possible to record the dynamic response of the respective sensors for *E*_GSH_ and H_2_O_2_ in plant cells, leaf samples or even whole seedlings using a microplate fluorescence reader. Importantly, genetic targeting of the sensors maintains subcellular specificity of the measurements across the measured tissues, even though the structures are not individually resolved. This approach is particularly well suited for measurements over several hours, can cover high numbers of samples, replicates and/or controls in parallel and can be expanded towards *in situ* biosensor multiplexing [21, 24, 25].

Here, we report protocols on how to use the Grx1-roGFP2 and roGFP2-Orp1 probes in both CLSM- and plate reader-based experiments for live monitoring of ROS-dependent redox changes in the cytosol. All approaches described here can be extended to measurements in other subcellular compartments, which would require appropriate targeting of the genetically-encoded probes.

## 2 Materials

### 2.1 Plant Material and Growth Conditions

1. Seeds of Arabidopsis lines constitutively expressing Grx1-roGFP2 or roGFP2-Orp1 in the cytosol.
2. Media plates for growing young seedlings: Agar plates with 0.5x Murashige and Skoog growth medium, 0.1% (w/v) sucrose, 0.05% (w/v) MES (pH = 5.8, KOH) and 0.8% (w/v) phytoagar (*see* **Note 1**)
3. Pots for growing plants in soil to rosette stage: Jiffy-7^®^-pellets (Jiffygroup, Oslo, Norway) (*see* **Note 1**).
4. Growth chamber capable of maintaining controlled conditions with a long day regime (16 h light, 100-120 μmol photons m^-2^ s^-1^, 19°C; 8 h dark, 17°C) with a relative humidity of 50%.

### 2.2 Assay Media and Stock Solutions

1. Imaging buffer: 10 mM MES, 10 mM MgCl_2_, 10 mM CaCl_2_, 5 mM KCl, pH = 5.8 supplemented with the different compounds for specific experiments.
2. 5-10 mM H_2_O_2_ and 5-10 mM DTT (dithiothreitol) dissolved in imaging buffer to achieve full oxidation or reduction of the sensors, respectively.
3. 100 mM Luminol (5-amino-2,3-dihydrophthalazine-1,4-dione) dissolved in DMSO (dimethyl sulfoxide).
4. 10 mg mL^-1^ horseradish peroxidase (HPR) dissolved in distilled water.
5. 1 mM flg22 peptide dissolved in distilled water.

### 2.3 Confocal Laser Scanning Microscopy and Perfusion Setup

1. Confocal microscope equipped with laser lines 405 nm and 488 nm to excite both roGFP2-based sensors and fluorescence detection in the 505 nm to 530 nm band. Here, a Zeiss LSM780 confocal microscope (Carl Zeiss Microscopy GmbH, Jena, Germany) equipped with a 25 mW Ar/ML-laser and a 30 mW 405 nm diode laser was used, but other confocal microscopes with laser lines of the same wavelengths are suitable, assuming sequential excitation is possible.
2. A 25x (NA 0.8) objective to image root regions or epidermal tissue areas or a 40x (NA 1.2) is for imaging single cells of the cotyledon, leaf or hypocotyl epidermis (*see* **Note 2**).
3. Slides and coverslips.
4. Featherweight forceps.
5. Adhesive rubber tape.
6. RC-22 perfusion chamber mounted on a P1 platform (Warner Instruments, Hamden CT) or similar.
7. Steel anchor harp with a 1.5 mm grid mesh to mount the seedling inside a perfusion chamber. The model SHD-22L/15 (Warner Instruments) was used for the RC-22 chamber.
8. Open 50 mL syringes connected to a VC-8M valve controller (Warner Instruments).
9. 1.5 mm polyethylene tubes (Warner Instruments).
10. Peristaltic pump (Ismatech, Wertheim, Germany) (**Fig. 2A**).

**Fig 2.**
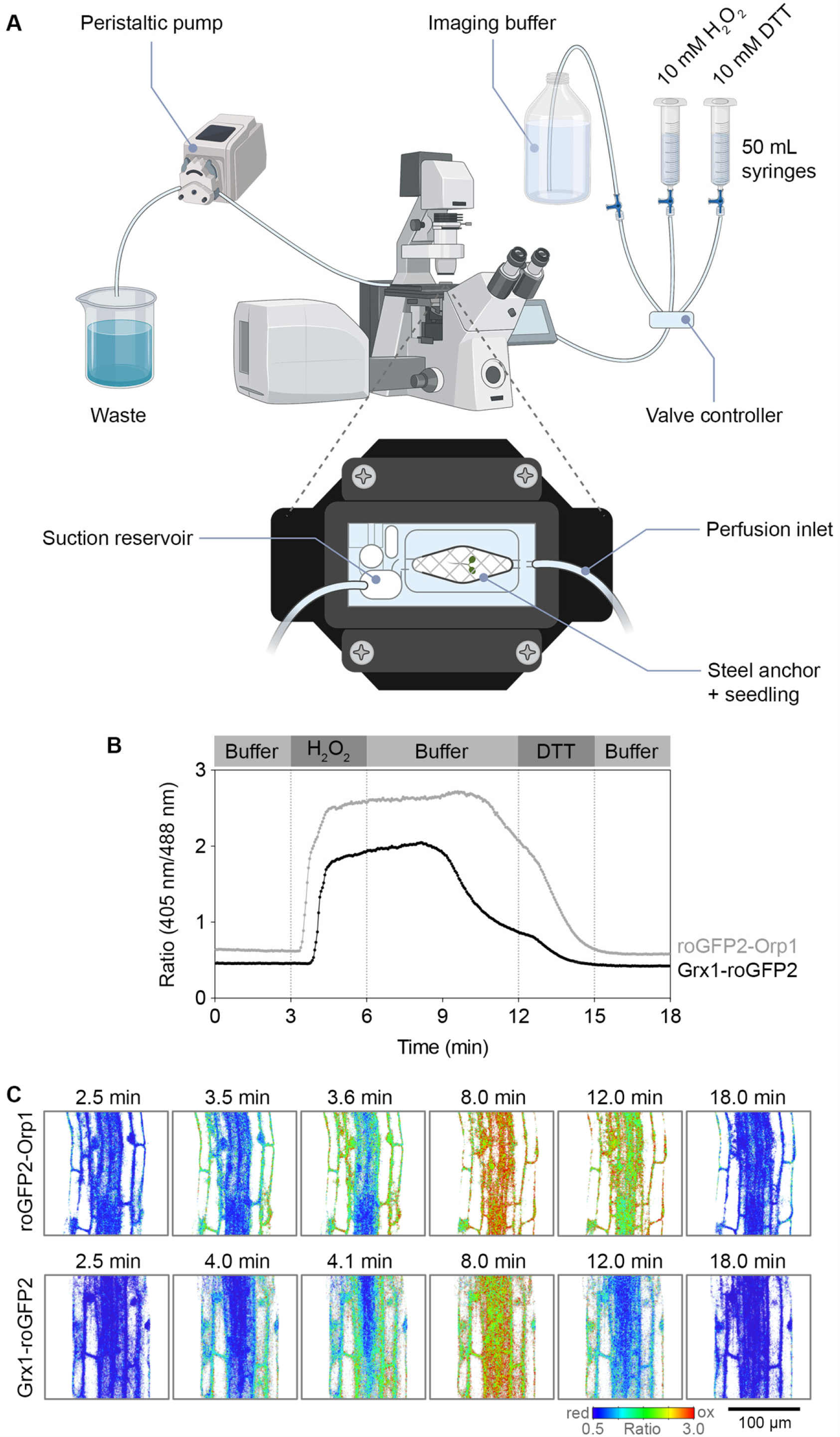
Live monitoring of roGFP2-based sensors using a perfusion system. (**A**) Scheme of the perfusion setup (made in ©BioRender - biorender.com). All treatments are provided in 50 mL syringes and fed into the chamber by gravity. The influx of each treatment solution is set by a central valve controller with a single output connected to the perfusion chamber. In the chamber, the plant tissue is held in place by a steel anchor and all excess media are continuously collected through suction generated by a peristaltic pump and discarded. (**B**) Calibration of roGFP2 probes in Arabidopsis roots. Root tissue of wild type plants expressing cytosolic Grx1-roGFP2 or roGFP2-Orp1 were treated as indicated in the plot, and the fluorescence ratio 405 nm/488 nm was recorded over time. (**C**) False-colored ratio images indicate the redox state of the sensors at different points during the time course.

### 2.4 Plate Reader Setup

1. Plate reader equipped with a monochromator and suitable filters for the two excitation maxima of both roGFP2-based sensors (e.g., 400 ± 5 nm and 480 ± 5 nm) and a 520 ± 5 nm emission filter, and a detector capable of detecting fluorescence and luminescence. Here, a CLARIOstar plate reader (BMG Labtech, Ortenberg, Germany) was used.
2. Polystyrene 96-well microtiter plates (Sarstedt, Nümbrecht, Germany) with transparent flat bottom.
3. Dissecting needle.

### 2.5 Image and Data Analysis Software

1. Redox Radio Analysis (RRA) software for MATLAB (The MathWorks, Natick, MA) (Fricker, 2016, https://markfricker.org/).
2. MARS Data Analysis Software (BMG Labtech).
3. MS Excel.
4. Statistical computing software such as R (https://www.r-project.org/) or GraphPad Prism (GraphPad Software, San Diego, CA) (https://www.graphpad.com/scientific-software/prism/).

## 3 Methods

The methods described below are based on the genetically encoded roGFP2 reporter constructs Grx1-roGFP2 and roGFP2-Orp1. Both probes have been characterized *in vitro* and *in vivo* and have been frequently used for live monitoring of *E*_GSH_ and H_2_O_2_, respectively. The biochemical and biophysical properties of the probes are summarized in **Table 1** (*see* **Note 3**).

**Table 1:** Characteristics of roGFP2-based sensors for the glutathione redox status (*E*_GSH_) and hydrogen peroxide (H_2_O_2_).

| Parameter | $E_{\text{GSH}}$ | $\text{H}_2\text{O}_2$ |
| --- | --- | --- |
| Sensor | Grx1-roGFP2 | roGFP2-Orp1 |
| Localization | Cytosol (*see Note 2*) |  |
| Promoter (*see Note 4*) | <sup>pro</sup>*UBQ10*<br>(from Arabidopsis) | <sup>pro</sup>*35S*<br>(from CaMV) |
| Plasmid backbone | pBinCM | pH2GW7 |
| Resistance in plants | Kanamycin | Hygromycin |
| Fluorescent protein | EGFP | EGFP |
| Operator/Sensor | Human glutaredoxin 1<br>(hGrx1) | Yeast glutathione peroxidase like 3<br>(Gpx3/Orp1) |
| Excitation maximum (**Fig. 1**) | 395 nm, 490 nm |  |
| Excitation wavelength<br>(for CLSM) | 405 nm, 488 nm |  |
| Excitation filter<br>(for plate reader) | $400 \pm 5$ nm, $480 \pm 5$ nm | |
| Emission maximum | 511 nm |  |
| Emission bandwidth<br>(for CLSM) | 505 to 530 nm |  |
| Emission filter<br>(for plate reader) | $520 \pm 5$ nm | |
| Ratio calculation | Excitation A-band / Excitation B-band<br>(**Fig. 1**; *see Note 6*) |  |
| Mid-point potential | -280 mV |  |
| Dynamic range<br>(purified sensor, *in vitro*) | $\sim 12$<br>390/480 nm | $\sim 8$<br>(pH > 6.5)<br>400/482 nm |
| Dynamic range<br>(in plants) | $\sim 5$ (CLSM,<br>405 nm/488 nm)<br>$\sim 3.5$ (Plate reader,<br>400 nm/480 nm) | $\sim 6.5$ (CLSM,<br>405 nm/488 nm)<br>$\sim 4.2$ (Plate reader,<br>400 nm/480 nm) |
| Linear range for measurements | -245 to -315 mV | N/A |
| Key references | [13–15, 22, 32] | [20, 33, 34] |

### 3.1 CLSM Methods

#### 3.1.1 Sample Mounting for Steady State Measurements

1. Prepare a microscope slide with two stripes of standard adhesive rubber tape to create a 180-250 µm spacer that will protect the sample from being squeezed by the coverslip (depending on the exact objective used this is typically #1.5; thickness 0.17 mm).
2. Screen Arabidopsis seedlings grown on vertically oriented agar plates for fluorescence under a stereo microscope equipped with the excitation/emission filters for GFP. Indicate the positive seedlings on the plate with a marker pen (*see* **Notes 4** and **5**).
3. Carefully take a seedling using featherweight forceps and place it on the microscope slide pre-loaded with a drop of imaging buffer.
4. Place a coverslip over the sample and secure it with two pieces of adhesive tape.
5. If the slide is not completely filled with imaging buffer, add additional buffer from the edge to fill the space between the slide and the coverslip.
6. Mount the slide on the microscope stage and secure it with stage clips.
7. Select a lens. The 40x lens (NA 1.2) is recommended to image whole epidermal cells, while the 25x (NA 0.8) is recommended to see entire root tips.
8. Illuminate the sample in transmission mode, locate and focus on the tissue to image.
9. Perform the measurements as indicated in section 3.1.3 or 3.1.4.

#### 3.1.2 Perfusion System Setup

1. Place a 22 × 40 mm coverslip (typically #1.5; thickness 0.17 mm depending on the exact objective used) on the bottom of the perfusion chamber and secure it with the side clamps (**Fig. 2A**).
2. Remove the piston from 50 mL syringe and fill each syringe with either 5-10 mM H_2_O_2_, 5-10 mM DTT, or imaging buffer. It is recommended to fill the syringes up to the same levels to start with the same flow. For this setup the gravity-dependent flow rate is 10 mL min^-1^, but may be modified by different tubing or restricting the flow.
3. Prime the tubes connected to the syringes carefully removing all air bubbles using the valve controller.
4. Place the steel anchor harp on a small petri dish, with the mesh facing upward, on a drop of water.
5. Carefully place a 4-5-day-old Arabidopsis seedling (pre-screened for fluorescence, *see* section 3.1.1) on top of the mesh using featherweight forceps (*see* **Note 7**). Avoid the area of interest to be imaged being too close to the mesh since its material is autofluorescent when illuminated at 405 nm.
6. Put a drop of imaging buffer into the perfusion chamber and insert the steel anchor harp. The sample should remain between the coverslip and the mesh.
7. Mount the chamber on the microscope stage and secure it with stage clips.
8. Start perfusing imaging buffer into the chamber and close the top of the chamber with a second coverslip.
9. Connect the suction tube to the suction reservoir and collect the excess volume with a peristaltic pump. Adjust the aspiration rate to be equal to the flow into the perfusion chamber.

#### 3.1.3 Elicitor-induced ROS Generation and Sensor Calibration

1. Incubate a 2-week-old Arabidopsis seedling (pre-screened for fluorescence, see section 3.1.1) in a 10 µM flg22 solution for 15 or 30 minutes.
2. Carefully transfer the seedling to a microscope slide between two stripes of adhesive tape (*see* section 3.1.1).
3. Carefully place a coverslip on top of the sample, without damaging the seedling.
4. Proceed as described in section 3.1.1, step (5) onwards.
5. To calibrate the sensor on the perfusion system, perfuse for 10 minutes 5-10 mM H_2_O_2_ and 5-10 mM DTT (*see* **Note 8**). We recommend perfusing imaging buffer for at least 10 minutes between treatments (**Fig. 2B-C**). To calibrate the sensor on steady state measurements, incubate the seedlings for 10 minutes in 5-10 mM H_2_O_2_ to fully oxidize or 5-10 mM DTT to fully reduce the sensor (*see* **Note 8**). After incubation in either H_2_O_2_ or DTT, place the seedlings carefully on a slide with a drop of the selected treatment solution for steady state measurements (**Fig. 3**). Root tips are best for imaging due to their permeability, but treatments on cotyledons or hypocotyl are possible (*see* **Note 8**).

**Fig 3.**
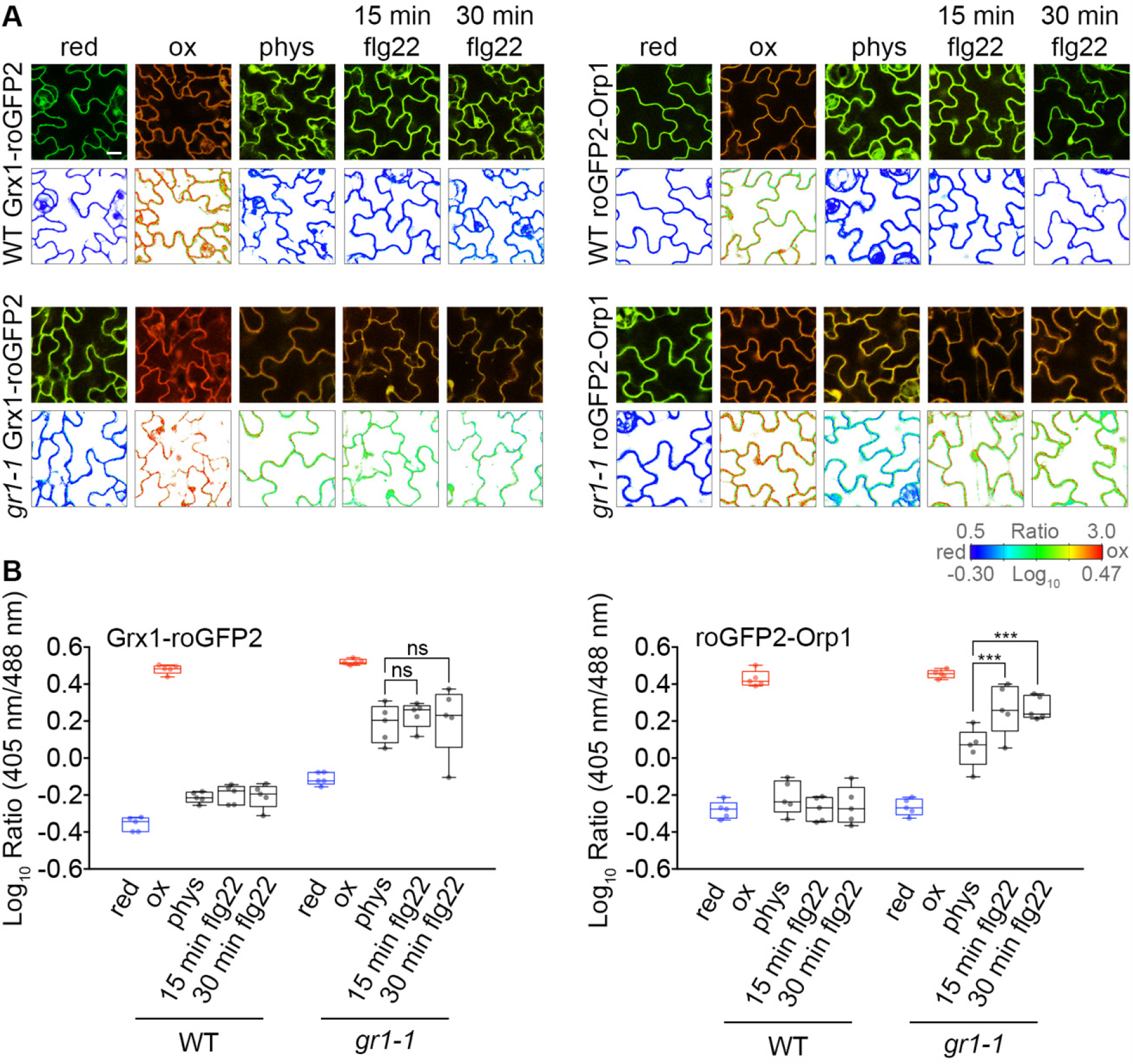
Cytosolic Grx1-roGFP2 and roGFP2-Orp1 respond to elicitor-induced oxidation in epidermal cells of *A. thaliana* leaves. True leaves of two-week-old Arabidopsis wild type plants (WT) and the GR1-deficient mutant *gr1-1* expressing Grx1-roGFP2 and roGFP2-Orp1 were treated with the bacterial elicitor flg22 (10 µM) for different time periods (15 and 30 min) to visualize the oxidative response. As control, leaves were treated with imaging buffer. For calibration, the samples were fully reduced (red) and fully oxidized (ox) with 10 mM DTT and 10 mM H_2_O_2_, respectively. (**A**) For confocal live cell imaging, both roGFP2-based sensors were excited at 405 and 488 nm and fluorescence collected from 508-530 nm. The upper panels are a merge between images from the 405 and 488 nm channels. The false-colored ratio images indicate the redox state of the sample (lower panels). Scale bars = 10 µm. (**B**) Quantitative analysis of flg22 induced oxidation in Arabidopsis leaves. Box plots indicate the log_10_ of the fluorescence ratios 405/488 nm of all measured samples expressing Grx1-roGFP2 or roGFP2-Orp1. For statistical analysis, a two-way ANOVA with Tukey’s multiple comparison test and a confidence level of *P* = 0.05 was conducted. Line = median, whiskers = min to max. *n* = 5.

#### 3.1.4 CLSM Settings

1. Set up two tracks for imaging roGFP2-based sensors with excitation at 405 nm and 488 nm, respectively. roGFP2 fluorescence is to be recorded in one channel between 508 and 530 nm for each track. An extra channel on the 405 nm track is used for detecting sample autofluorescence between 430 and 470 nm, while an extra channel on the 488 nm track can be used to collect chlorophyll autofluorescence in green tissues.
2. Select line switching as scan mode for the two tracks and an averaging of maximum two scans. Set the pixel dwell time to 1.27 µs/pixel and the frame size to 512 × 512 pixels with a data depth of 12 bits. These settings will limit the required time for imaging to about 2.46 s per frame, which allows monitoring fast redox changes during perfusion treatments.
3. Adjust the master gain to the same value (within the linear range of the detector at a maximal value of 800) for the channels detecting roGFP2 and autofluorescence excited at 405 nm. For quantitative imaging, it is crucial, that neither the laser power, nor the detector gain of these channels is changed during an experiment. For the transmitted light channel and chlorophyll autofluorescence, the gain can be adjusted independently.
4. Start the ‘live’ preview scan of the sample fluorescence with the range indicator lookup table for image display. Adjust the power of the 488 nm laser until a clear and structured fluorescent signal is visible from the sensor. Adjust gain and offset to set the image into the dynamic range: The master gain for the roGFP channels (405 nm and 488 nm excitation) and the autofluorescence channel (405 nm excitation) must be kept identical over the course of the experiment to allow quantitative comparison. Increase the power of the 405 nm to double the power output of the 488 nm laser (e.g., 2% power for 488 nm; 4% power for 405 nm) (*see* **Note 9**).
5. After adjustment of the imaging settings it is crucial to keep them unchanged between samples. Any modification will affect the resulting ratios and make comparison of fluorescence ratios between samples impossible.
6. Take an image of an analogous tissue region of a wild-type plant with your final settings in order to determine the bleed-through of autofluorescence excited at 405 nm to your GFP emission channel.

#### 3.1.5 Data Collection and Analysis

1. Steady-state images or time series are recorded by the microscope software as Zeiss LSM files (*.lsm) or other formats. A minimum of 12 images is recommended to compare genotypes or treatments.
2. Either use the MatLab-based Redox Radio Analysis (RRA) software [30] tailored to the analysis of data of genetically encoded biosensors, or other image analysis software such as e.g., Image J.
3. Subtract the background from each image.
4. Correct the roGFP2 emission channel at 405 nm excitation for bleed-through of autofluorescence excited at 405 nm: Use images taken on wild type samples (*see* section 3.1.4 step (6)) to determine the factor between the autofluorescence measured between 430 and 470 nm and the bleed-through to the roGFP emission channel. Subtract the autofluorescence measured for each image multiplied by this determined factor from the roGFP emission at 405 nm excitation. More detailed information is available in the RRA handbook (https://markfricker.org/) (*see* **Note 10**).
5. Calculate the ratio between the corrected roGFP2 emission at 405 nm excitation and the roGFP2 emission at 488 nm excitation for each pixel in the image (*see* **Note 11)**.
6. Generate a ratio image by using the values measured during calibration as minimum (0% sensor oxidation) and maximum (100% sensor oxidation) values of the color scale.
7. Calculate the average 405/488 ratio for each image.
8. Log_10_-transform ratio values (*see* **Note 12**) and do statistical analysis of differences between treatments or genetic backgrounds.
9. Fluorescence ratios may be converted to the degree of oxidation of roGFP2 and/or *E*_GSH_ in mV. For further details *see* [14, 23, 26].

### 3.2 Plate Reader Methods

#### 3.2.1 Sample Mounting of Whole Seedlings

1. Pre-screen the vertically grown seedlings for fluorescence (*see* section 3.1.1)
2. Fill a 10 cm petri dish with 10 mL imaging buffer and pipette 200 µL imaging buffer in all wells required for the experiment (*see* **Note 13**).
3. Use a modified dissecting needle with its tip being bent to form a hook, to carefully pick up a single seedling at the hypocotyl and transfer it into the imaging buffer in the petri dish. This action will discharge the static electricity on the seedlings, which makes it easier to place them in the 96-well plate. Repeat the process to pool 3-4 seedlings per well and set up at least 4-6 replicates for each treatment (*see* **Notes 14** and **15**).
4. To calibrate the sensor at the beginning of the experiment, transfer seedlings to four wells filled with either 5-10 mM H_2_O_2_ or 5-10 mM DTT in imaging buffer, to fully oxidize or reduce the sensor, respectively. Alternatively, calibrations can be done on each single well at the end of the experiment by replacing the buffer first with 5-10 mM DTT and subsequently with 5-10 mM H_2_O_2_. Fluorescence readings need to be taken for samples immersed in DTT and H_2_O_2_, respectively.

#### 3.2.2 Sample Mounting of Leaf Discs

1. For leaf discs, 1-2 discs of 8 mm diameter are cut with a cork borer from a fully expanded leaf of 4-week-old plants. Since the cut is made from the adaxial side, it is recommended to use a flat surface as support from the abaxial side of the leaf. For instance, a piece of cardboard may be used.
2. Transfer a maximum of 8 discs per well on a 6 well-plate pre-loaded with 2 mL imaging buffer using a pair of featherweight forceps. Float the discs with the abaxial side down and incubate them in the dark overnight to minimize sensor oxidation due to the wounding stress. After incubation, discard the buffer and refill with fresh buffer in order to remove any leaked ions after the cut. Transfer each leaf disc to a single well on a 96 well-plate pre-loaded with 200 µL imaging buffer.
3. Push the disc to the bottom of the well using a second pair of forceps. Since the well (7 mm) is slightly smaller than the disc (8 mm), the disc should not move from the bottom (*see* **Note 16)**.
4. As mentioned in step (4), control discs from the same genetic background without sensor expression should be used for background and autofluorescence subtraction.
5. Calibration of the sensor should be performed as indicated in step (4).

#### 3.2.3 Settings for Spectral Measurements

1. Place the 96 well-plate with the seedlings or leaf discs in the plate reader.
2. Define the layout of the samples in the plate reader software.
3. Run an excitation spectrum scan, from 350 nm to 495 nm while collecting the fluorescence with the 520 ± 5 nm emission filter (*see* **Note 17**).
4. Set the gain and focal height. Use 480 nm excitation, since it is close to excitation maximum of roGFP2 yielding a more intense fluorescence signal. Gain and focal height should be measured in at least 6 different wells and the average value should be used (*see* **Note 18**). It is recommended to adjust the gain to reach 50% of the maximum detectable fluorescence in order to avoid saturation of the detector, which would invalidate the measurement. As a reference, use samples incubated in 10 mM DTT.
5. Set a high number of flashes per well (> 30). This will create an average value for each excitation wavelength with improved signal-to-noise, which will result in smooth fluorescence spectra.
6. If available, define the recording as ‘orbital averaging’ with flashes exciting the sample along a circle with no less than 3 mm diameter. Averaging the fluorescence of different areas of the same sample will increase the reliability of the measurements. This is particularly important when the plant tissue does not cover the full well area, as typically the case for seedlings.

#### 3.2.4 Settings for Ratiometric Time Course Measurements

1. Place the 96 well-plate with the seedlings or leaf discs in the plate reader.
2. Define the layout of the samples in the software.
3. If auto-injection is required, prepare the treatment stock solutions and prime the injectors within the plate reader. It is recommended to prepare no less than 5 mL since the pump cylinder of the auto-injection pump holds 500 µL.
4. Set the excitation wavelengths for the two channels to 400 ± 5 nm and 480 ± 5 nm, respectively. In both cases, the emission is set to 520 ± 5 nm (*see* **Note 19**).
5. Set the gain and focal height as mentioned in section 3.2.2 step (4). Subsequently, follow the same routine to set the gain for excitation at 405 nm on a sample incubated in 5-10 mM H_2_O_2._
6. Adjust the number of flashes per well and orbital averaging depending on the required speed of measurement (*see* section 3.2.2 steps (5) and (6)). To document fast reactions, a lower number of flashes is recommended. In contrast, measures for the calibration or steady states for roGFP2 fluorescence might benefit from increased averaging, which is achieved by more flashes per well.

#### 3.2.5 Elicitor-induced ROS Generation

1. Adjust the layout of the experiment on the plate reader software.
2. Prepare an assay solution of 20 µM flg22 from the stock.
3. Transfer the leaf discs from four-week-old Arabidopsis plants, expressing any of the roGFP2-based sensors to the wells of a 96 well-plate pre-loaded with 50 µL imaging buffer (*see* **Note 20**).
4. Add 50 µL of the assay solution to each well and start the measurement immediately.
5. Record the fluorescence of the channels as described in section 3.2.3 for 2 h (**Fig. 4**).
6. To measure extracellular ROS production, prepare the assay solution with 200 µM Luminol, 20 µg/ml HRP, and 20 µM flg22 from the respective stock solutions (*see* section 2.3).
7. Transfer the samples as indicated in steps (3) and (4).
8. Record the luminescence (**Fig. 4**).

**Fig 4.**
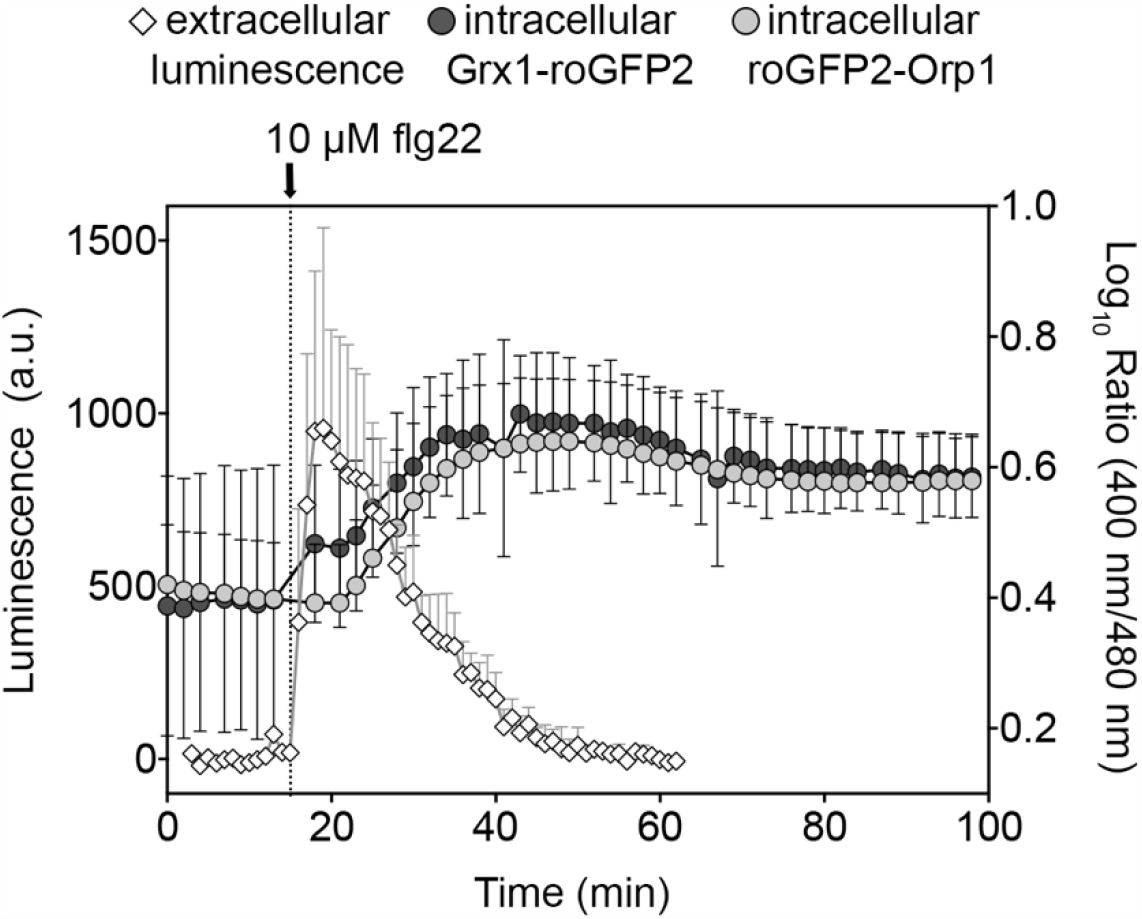
Time-resolved measurement of the extracellular and intracellular oxidative burst after elicitor-treatment of *A. thaliana* leaf discs. The combined graph shows the extracellular luminescence after reaction of luminol with ROS in the apoplast (open diamonds) and the intracellular response of Grx1-roGFP2 (dark grey circles) and roGFP2-Orp1 (light grey circles) after the treatment of *gr1-1* leaf discs with 10 µM flg22 (arrow). Data indicates the mean value ± SD of the log_10_ of the 400 nm/480 nm fluorescence ratios collected from the entire tissue in the well. All measurements were done on a plate reader. For luminescence, *n* = 3, for fluorescence, *n* = 5.

#### 3.2.6 Data Collection and Analysis

1. For each comparison (genotype or treatment) a minimum of 8 replicates of seedling pools or leaf discs is recommended. Depending on the layout, these can be measured on the same run side by side. For non-fluorescent control samples, 5 replicates are sufficient.
2. Export the raw data to a spreadsheet using the data analysis software of the plate reader.
3. For processing of the fluorescence data, average the values of the non-fluorescent samples and subtract the average value from the fluorescence recorded for samples expressing the sensors. Apply this for each condition tested (e.g., control and treatment). For each time point, divide the corrected fluorescence values of channel 1 (400 nm excitation) by the corrected fluorescence values from channel 2 (480 nm excitation) (*see* section 3.2.3).
4. The dynamic range of the sensor can be estimated as the value obtained from the ratio values for fully oxidized and fully reduced samples.

## 4 Notes

1. The age of Arabidopsis plants to use varies depending on the research question and requires careful individual consideration of technical and biological issues. 2-5-day-old seedlings grown vertically on plates under axenic conditions are particularly suitable for CLSM imaging of roGFP2-based biosensors. The small size at this developmental stage allows placement of whole seedlings in a perfusion chamber and enables whole seedling imaging. However, there may be other points to consider, such as the non-uniform expression pattern of *FLS2* coding for the flagellin receptor, which would be critical for observation of ROS production induced by the elicitor flg22 [27, 28]. To obtain 4-week old plants is recommended to grow them on Jiffy-7^®^-pellets.
2. To image the sensor targeted to subcellular compartments, such as mitochondria, a 40x (NA 1.2) water immersion objective is recommended [31]. Oil immersion lenses with NA 1.4 can provide results at high resolution for tissue layers just below the coverslip. To image whole seedlings, use a 10x and take various images that can be stitched using image editing software like Gimp, Adobe Photoshop or the ZEN software from Zeiss. Lower magnification lenses help avoiding photodamage.
3. Both sensors have also been targeted to mitochondria by fusion with the transit peptide sequence either from Serine Hydroxymethyltransferase (SHMT; for roGFP2-Grx1; [29]) or from the *Nicotiana plumbaginifolia* ATP synthase ß-subunit of (for roGFP2-Orp1; [21]), or to plastids by cloning the Transketolase transit peptide (TKTP) to the N-termini of the sensors [13]). All lines expressing these constructs are available upon request. For measurements on mitochondria, *see* [30].
4. Screening by fluorescence for transformants using a fluorescence stereomicroscope is particularly useful if resistance markers that are typically used for selection are present in the background already. In addition, screening or control for fluorescent plant materials under a stereomicroscope enables a first evaluation of the expression levels and potential inhomogeneity of the expression. Constitutive expression driven by the *CaMV 35S* or *UBQ10* promoter may lead to silencing of the transgene as apparent from a decrease of fluorescence over generations and patchy expression of the sensor across a seedling.
5. Some reporter lines, like e.g. lines with roGFP targeted to mitochondria or roGFP expressed in some mutant backgrounds can have a lower fluorescence compared to WT plants with cytosolic roGFP2 constructs driven by a strong promoter. This low fluorescence may be difficult to recognize under the stereomicroscope and thus selection of suitable reporter lines requires particular care.
6. The fold change in fluorescence intensities along the entire excitation spectrum is not constant (**Fig. 1B**). Therefore, the achievable dynamic range depends on the chosen wavelength combination. For selection of the most suitable wavelength combination further parameters like absolute fluorescence intensities and signal-to-noise need to be taken into account.
7. Mount the seedling with the shoot oriented towards the inlet. This will decrease the risk of unintended movements or the root tip.
8. The appropriate concentration of DTT and H_2_O_2_ for the calibrations may vary with different tissues and organs and needs to be determined first. If no complete reduction or oxidation is achieved, a 10 min vacuum infiltration is recommended (for calibration purposes only) or increase the concentrations up to 50 mM. For reference values for the dynamic range *see* **Table 1**.
9. If the sample fluorescence is too low, open the pinhole rather than increasing the laser power. This will minimize the risk of photodamage. To check if chosen settings are suitable for quantitative imaging, the *in vivo* calibration of the sensor in the respective plant material should be done first. Neither in the fully reduced nor the fully oxidized state many pixels should be saturated. After settings are chosen for quantitative imaging, pixel saturation can only be controlled through pinhole adjustment. Initial pinhole size should depend on the size of the imaged object (e.g., plastids vs. whole cells). Changing the pinhole diameter will also change the volume of sampling and therefore the thickness of the cell layer imaged.
10. Load each *.lsm file into the RRA software and select a probe MatLab file with all the pre-set parameters for the analysis of each sensor. These parameters include among others, the sensor midpoint redox potential, and identification of the different channels. The range values (R_min_/R_max_) for the false colored images of the 405 nm/488 nm fluorescence ratio need to be set according to the values obtained by the *in vivo* calibration. Follow the software interface to subtract the autofluorescence, align the channels if needed and calculate the fluorescence ratio. The software will export the ratio data of all batched samples to an MS Excel file and the false colored images as TIFF files. Express the dynamic range of the sensor as the value obtained from ratio_ox_/ratio_red_.
11. With purified recombinant protein the maximum achievable spectroscopic dynamic range with 405 nm and 488 nm excitation is about 9-fold for free roGFP2 and Grx1-roGFP2 and about 6-fold for roGFP2-Orp1 [21, 23]. In live cells, the dynamic range is often slightly lower, typically between 3- and 6-fold. The exact value largely depends on the quality of the background and autofluorescence correction. Since the spectroscopic dynamic range at a given excitation wavelength combination is characteristic for a given biosensor and independent of instrument settings, it provides a control parameter for *in vivo* measurements that cannot exceed the corresponding *in vitro* value of the purified sensor protein. For further details see [23, 26].
12. The correlation between the absolute value of the mean and the variance in ratio data implies that higher ratios have bigger standard deviations. The unequal variance can be corrected for by log_10_ transformation, which will convert the skewed data distribution to a normal distribution.
13. Lower amounts of buffer can be also used, but the minimum recommended is 100 µL, to evenly cover the bottom of the well and to avoid evaporation during the run.
14. For plate reader measurements in 96-well plates, use 7 day-old vertically-grown seedlings and a pool of 3-4 seedlings per well to obtain a good signal while limiting the time for sample preparation. To measure the sensor fluorescence in mature leaves, leaf discs or leaf rolls from 4-week-old plants can be used. In all cases, sufficient material is needed to cover the well area.
15. It is recommended to measure at least 4–6 wells and average the calculated ratios to account for variability between individual wells. Before ratiometric analysis, all datasets should be pre-screened for any obvious technical inconsistencies. These might be fluorescence ratios far outside the plausible and calibrated spectroscopic dynamic range (*see* **Note 11**), a pronounced drift of both channels in the same direction during the course of an experiment, or lack of converse changes of the two fluorescence channels. Such problems might be caused could by movement of samples in the wells or technical issues of the plate reader detection. To avoid including erroneous datasets, it is important to check that the fluorescence of the individual channels responds in the expected opposing manner and that all other parameters are fulfilled. Replicates that do not meet these criteria should be treated as outliers and not be considered for further analysis.
16. If leaf discs are treated with H_2_O_2_ for extended periods, the cells might lose their turgor and leaf discs will float atop the medium. This will lead to loss of proper focal adjustment and hence affect the fluorescence measurements.
17. The upper wavelength for the excitation spectrum is defined by the choice of the dichroic mirror and the emission wavelength. If spectra beyond the nominal emission maximum of the fluorophore are to be collected, appropriate dichroic mirrors and longer emission wavelengths need to be used.
18. Leaf discs placed at the bottom of the wells give uniform measurements of focal height while pools of seedlings can give a range of focal heights. In the latter case, it is recommended to measure at least 6 wells and average the height.
19. Many plant tissues show increasing autofluorescence when moving to wavelengths lower than about 400 nm, but autofluorescence can start to increase also above 400 nm. If the signal-to-noise ratio is too low, it is recommended to use a longer wavelength for excitation depending on the available filters, e.g., 410 nm.
20. In order to detect elicitor-induced ROS bursts in the cytosol via the local glutathione redox potential, we recommend using *gr1-1* mutants to achieve higher sensitivity and better resolution.

## Acknowledgements

Support by the Deutsche Forschungsgemeinschaft through the Research Training Group GRK2064 (to AJM, MS and SJM-S) and through the priority program SPP1710 (to AJM and MS) is gratefully acknowledged. We thank our former lab members Thomas Nietzel, Stephan Wagner and Philippe Fuchs for their seminal work in establishing protocols and analysis routines for live redox imaging and plate reader assays.

## Notes

### Competing Interest Statement

The authors have declared no competing interest.

